# *Caenorhabditis elegans* BMP Transcriptional Program Implicates Collagen Genes in Body Size Regulation

**DOI:** 10.1101/108225

**Authors:** Uday Madaan, Edlira Yzeiraj, Michael Meade, Christine A. Rushlow, Cathy Savage-Dunn

**Affiliations:** Department of Biology, Queens College, and the Graduate Center, CUNY, Flushing, NY, USA; Department of Biology, New York University, New York, NY, USA

## Abstract

Body size is a tightly regulated phenotype in metazoans that is dependent on both intrinsic and extrinsic factors. While signaling pathways such as insulin, Hippo, and myostatin are known to control organ and body size, the downstream effectors that mediate their effects are still poorly understood. In the nematode *C. elegans*, a Bone Morphogenetic Protein (BMP)-related signaling pathway is the major regulator of growth and body size. DBL-1, the BMP-related ligand, is secreted by neurons and body wall muscle, and acts as a dose-dependent regulator of body size. We investigated the transcriptional network through which the DBL-1/BMP pathway regulates body size and identified cuticle collagen genes as major effectors of growth control. Here we demonstrate that cuticle collagen genes can act as positive regulators (*col-41*), dose-sensitive regulators (*rol-6*), and negative regulators (*col-141, col-142*) of body size. Moreover, we show requirement of DBL-1/BMP signaling for stage-specific expression of cuticle collagen genes. We used chromatin immunoprecipitation followed by high throughput sequencing (ChIP-Seq) and electrophoretic mobility shift assays to show that the Smad signal transducers directly associate with conserved Smad binding elements in regulatory regions of *col-141* and *col-142*, but not of *col-41.* Hence, cuticle collagen genes are directly and indirectly regulated via the DBL-1/BMP pathway. These results provide the first direct regulatory link between this conserved signaling pathway and the collagen genes that act as its downstream effectors in body size regulation. Since collagen mutations and misregulation are implicated in numerous human genetic disorders and injury sequelae, understanding how collagen gene expression is regulated has broad implications.

**Author Summary:** Body size in humans and other animals is determined by the combined influence of genetic and environmental factors. Failure to regulate growth and body size appropriately can lead to a variety of functional impairments and reduced fitness. Progress has been made in identifying genetic determinants of body size, but these have not often been connected into functional pathways. In the nematode model *Caenorhabditis elegans,* single gene mutations in the BMP signaling pathway have profound effects on body size. Here we have elucidated the BMP transcriptional network and identified cuticle collagen genes as downstream effectors of body size regulation through the BMP pathway. Collagens play diverse roles in biology; mutations are often associated with rare heritable diseases such as osteogenesis imperfecta and Ehlers-Danlos syndrome. Our work thus connects a conserved signaling pathway with its critical downstream effectors, advancing insight into how body size is specified.

## Introduction

The Transforming Growth Factor-beta (TGF-β) superfamily encompasses more than 30 ligands including Bone Morphogenetic Proteins (BMPs), Activin, and Nodal. BMPs play essential roles in development and have been studied in a variety of contexts [1]. The roles of BMPs in growth and body size regulation, however, are relatively unexplored. This deficit is partly due to body size being expressed as a complex trait with continuous variation. The existence of monogenic variants with discrete body size changes provides us with an entry point into growth and size regulating mechanisms. We have capitalized on the major visible effect of BMP signaling on growth regulation in the nematode *Caenorhabditis elegans* to uncover transcriptional targets acting as effectors of body size regulation.

BMPs are highly conserved among animal species along with the components of their signaling pathways. BMP homologs are present in Drosophila (e.g. Decapentaplegic (Dpp)) and *C. elegans* (e.g. Dpp/BMP-like-1 (DBL-1)) [1], genetically tractable organisms in which Smads, intracellular components of the TGF-β signaling pathway, were first identified [2, 3]. Receptor-regulated Smads (R-Smads) are directly phosphorylated by the TGF-β receptors on the C-terminus. This allows formation of the heterotrimeric Smad complex with co-Smads that accumulates in the nucleus to regulate target genes of the pathway [4–14]. Smads are known to bind a 4bp GTCT Smad Binding Element (SBE); furthermore, R-Smads for BMP ligands associate with GC-rich sequences (GC-SBE) [15–17]. While much detailed knowledge has been obtained in studies of a few direct Smad target genes, fewer studies have addressed the direct and indirect transcriptional programs required to mediate biological functions in intact organisms. In *C. elegans*, the DBL-1/BMP signaling pathway is the major regulator of growth and body size. DBL-1 ligand is secreted by neurons and body wall muscle, and is necessary for body size regulation among other developmental processes [18]. The small body size phenotype is the result of a reduction in cell size rather than cell number. Previous work has pinpointed the hypodermis, the outermost multinucleated epithelium, as the main target tissue of DBL-1 signaling. In *dbl-1* mutants, different tissue sizes are reduced to different extents, with hypodermal tissue reduced in proportion to body size [19]. Furthermore, we have previously shown by tissue-specific expression of SMA-3 (R-Smad) in a *sma-3* mutant background that activation of the DBL-1 pathway in the hypodermis is necessary and sufficient for normal body size [20]. Similar conclusions have been drawn in experiments with the DBL-1 receptors [21, 22]. To identify the transcriptional targets of the DBL-1 pathway that may function in body size regulation, we performed microarray analysis on *dbl-1* mutants [23]. The functions of putative target genes were analyzed by RNA interference (RNAi) knockdowns, leading us to concentrate on a group of cuticle collagen genes.

The cuticle serves as the exoskeleton of the worm and its main component is collagen, encoded by more than 170 genes in the cuticular collagen multigene family [24]. The cuticle is synthesized and secreted by the underlying hypodermis, and polymerizes on the external surface forming the exoskeleton. At the end of each larval stage, the cuticle molts revealing a newly formed cuticle underneath. Some cuticle collagen genes are expressed in specific larval stages while others are expressed in each cuticle synthesis period [25]. In spite of extensive study, only a few of the cuticle collagen genes have been described to have visible mutant phenotypes; these abnormalities include body morphology defects such as DumPY (Dpy), ROLler (Rol), BLIster (Bli), Squat (Sqt), and LONg (Lon) [26]. We reasoned that altered expression of cuticle collagen genes could contribute to the small body size phenotype of DBL-1 pathway mutants.

We show here that cuticle collagen genes are direct and indirect transcriptional targets of the DBL-1 pathway. We demonstrate SMA-4 (co-Smad) binding in the intergenic region between *col-141* and *col-142*. This result is the first demonstration that the DBL-1 pathway Smads bind DNA *in vitro,* further validating the functional conservation of Smads in the nematode model. We find that other collagen genes are likely indirect targets of the DBL-1 pathway. Moreover, we observe loss of stage-specific expression of cuticle collagens in the absence of DBL-1 signaling. Lastly, we demonstrate through collagen loss-of-function mutants, RNAi, and overexpression studies that cuticle collagens are the effectors of body size regulation by the DBL-1 pathway.

## Results

### Cuticle collagen genes are transcriptional targets of the DBL-1 pathway and are required for normal body size

We sought to connect the DBL-1 transcriptional program to its downstream effectors of body size regulation. In previous work, we identified five putative target genes with body size phenotypes [23], but all of these genes encode signaling molecules or transcription factors, so they are not the downstream effectors of growth control. It has been proposed that changes in hypodermal polyploidy contribute to the reduced growth of *dbl-1* mutant adults [27], although these changes cannot account for differences in size during larval stages. We tested the functions of putative target genes encoding DNA licensing factors and cyclins that could regulate polyploidization, but knockdown of these genes by RNAi did not produce body size phenotypes (S1 Fig). We next turned our attention to four cuticle collagen genes that we identified as putative transcriptional targets of the DBL-1 pathway. We performed qRT-PCR to verify transcriptional changes in cuticle collagen genes. We compared expression in *dbl-1* and *sma-3* mutants to wild-type controls at the second larval (L2) stage, the same stage in which the microarray analysis had been conducted. In absence of DBL-1 signaling, *rol-6* and *col-41* are down regulated while *col-141* and *col-142* are up-regulated (Table 1), in agreement with the microarray studies.

**Table 1.** **Cuticle collagen genes are transcriptional targets of the DBL-1 pathway**

| Gene | L2 Stage |  |  | Adult Stage |
| --- | --- | --- | --- | --- |
|  | *dbl-1*<br>Microarray<br>Fold Change | *dbl-1*<br>qRT-PCR<br>Fold Change | *sma-3*<br>qRT-PCR<br>Fold Change | *sma-3*<br>qRT-PCR<br>Fold Change |
| *rol-6* | -2.15 | -1.3 | ND | 3.6 |
| *col-41* | -2.64 | -1.2 | -1.4 | 1.6 |
| *col-141* | 3.19 | 1.6 | 1.7 | -1.7 |
| *col-142* | 2.65 | ND | 1.5 | -1.2 |

Microarray results are reproduced from [23]. qRT-PCR results are from representative trials.
ND: Not determined.

We hypothesized that regulated expression of these cuticle collagen genes contributes to body size regulation. An alternative hypothesis is that the altered expression of cuticle collagen genes is a response to, rather than a cause of, the altered size of *dbl-1* mutants [28]. To address the role of these cuticle collagen genes in body size regulation, we performed RNAi of *col-41, rol-6,* and *col-141* and measured body length at different time points during larval and adult growth (Fig 1A). RNAi of *col-41* and *rol-6* led to a decrease in body size at all stages. Conversely, *col-141* RNAi led to a transient increase in body size in the larval stages, suggesting that cuticle collagens can act as negative as well as positive regulators of body size. We verified our results using an available loss-of-function mutation in *rol-*6. This *rol-6(lf)* mutant also has a small body size, as illustrated by body length measured at the L4 stage (Fig 1B).

**Fig 1.**
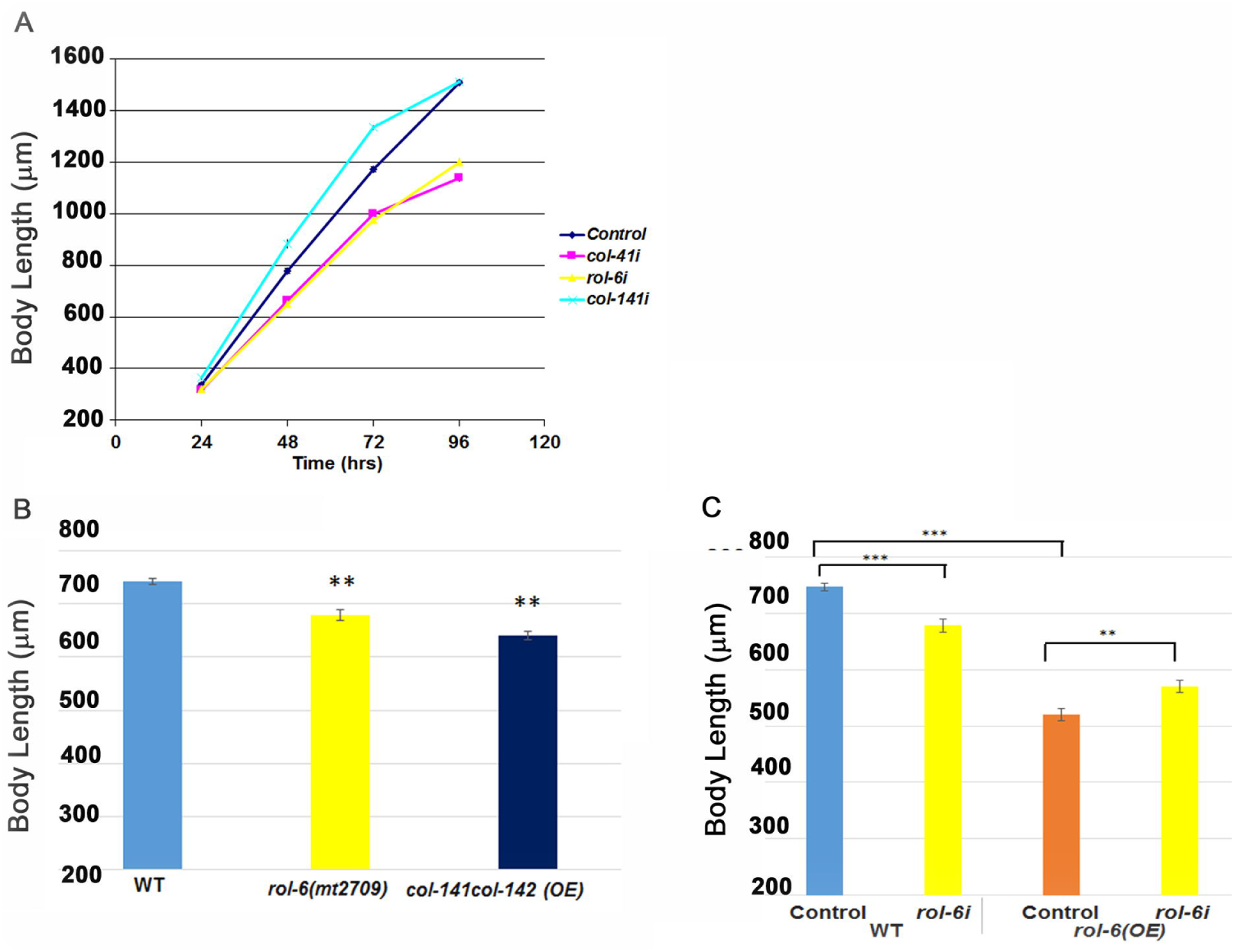
**Cuticle collagen gene inhibition and overexpression leads to body size phenotypes. A.** RNAi inhibition of *col-41*, *rol-6*, and *col-141* led to changes in body size. Animals were measured at indicated time points following embryo collection. **B**. Comparing body size phenotype of *rol-6(mt2709)* mutant and overexpressing *col-141col-142* worms relative to wild type. Body length was measured at the L4 stage. An independently generated transgenic line gave equivalent results. **C**. Overexpression of *rol-6* leads to smaller body size. *rol-6* RNAi in the *rol-6* overexpression line alleviates small body size. Body length was measured at the L4 stage. An independently generated transgenic line gave equivalent results. OE-overexpression; Control-Empty Vector. Error bars show standard error. * P < 0.05, ** P < 0.005, ***P < 0.0005.

If cuticle collagen expression is truly instructive rather than permissive for body size regulation, we would expect that overexpression of respective collagens should also lead to body size changes. Therefore, we performed overexpression of *col-141* and *col-142,* by increasing copy number in transgenics carrying a genomic fragment that contains both genes, which are adjacent in the genome. Overexpression of *col-141* and *col-142* resulted in decreased body size relative to wild-type worms (Fig 1B), confirming these genes as negative regulators of growth. We performed a similar experiment to overexpress *rol-*6. Surprisingly, overexpression of *rol-6* led to a decrease, rather than an increase, in body size (Fig 1C). To confirm that the observed phenotypes were due to *rol-6* overexpression, rather than an artifact of the transgene, we performed *rol-6* RNAi on the overexpression lines to reduce the level of overexpression. This RNAi inhibition of *rol-6* expression ameliorated the decreased body size in the overexpressing lines (Fig 1C). These results suggest that *rol-6* is dose-sensitive and must be expressed in a particular range; otherwise aberrant body size and morphological defects are observed.

### SMA-3 associates with Smad Binding Elements in the intergenic region between *col-141* and *col-142*

To identify which transcriptional targets of DBL-1 signaling were potentially directly regulated by Smad binding, genome-wide ChIP-seq was performed using a strain expressing functional GFP∷SMA-3 from the endogenous *sma-3* promoter, in a *sma-3(wk30)* background. Using the Galaxy software [29–31], we extracted genomic sequences from the coordinates obtained via ChIP-seq. MEME-ChIP software [32] was used to discover enriched motifs and generate logo plots. The top two enriched motifs were the CTGC<u>**GTCT**</u> SBE and GAGA (Fig 2A). The presence of GTCT as an enriched motif indicates an increased presence of SMA-3 at the indicated association sites. Among the cuticle collagen genes that we established as transcriptional targets, the intergenic region between *col-141* and *col-142* had strong Smad binding (Fig 2B). No additional SMA-3 recruitment sites were identified in this genomic region, suggesting that the intergenic region serves to regulate both of these genes. We therefore chose to study this region, which we will refer to hereafter as *col-p.* Further analysis of *col-p* revealed multiple SBEs that are conserved between three related nematode species (Fig 3A).

**Fig 2.**
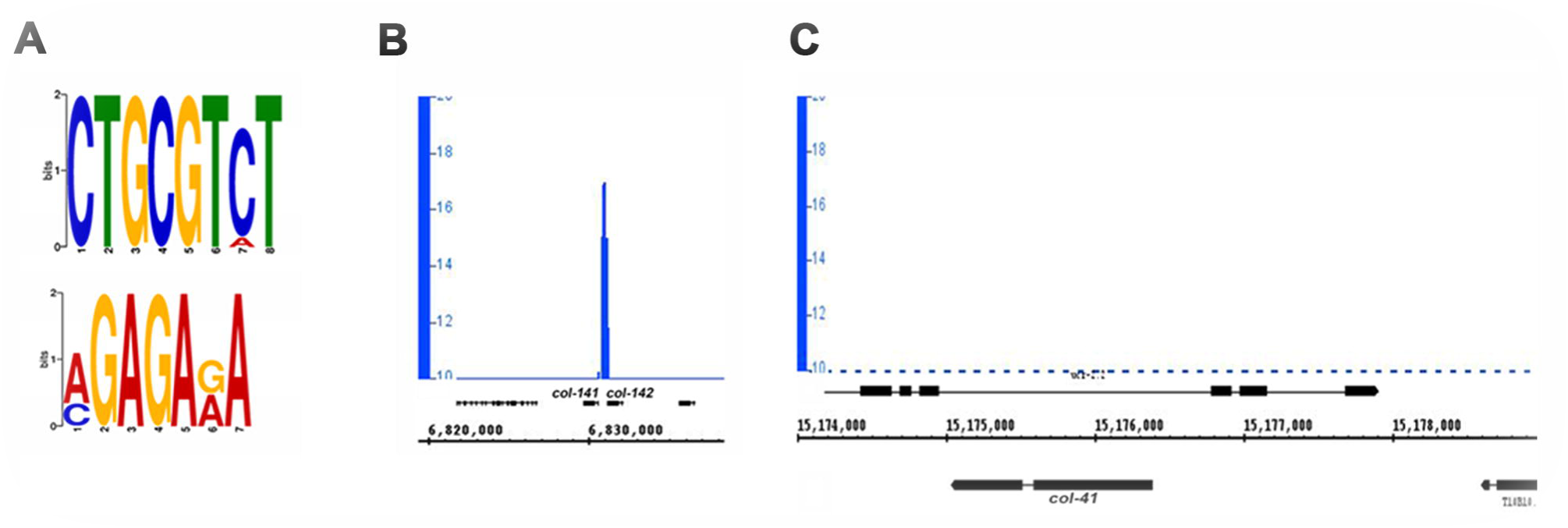
**SMA-3 associates with Smad Binding Elements in the intergenic region between *col-141* and *col-142 (col-p)*. A**. SMA-3 ChIP sequencing revealed SBE as highly enriched motif via MEME-ChIP motif discovery analysis. **B**. SMA-3 associates with the intergenic region between *col-141* and *col-142 (col-p)*.

**Fig 3.**
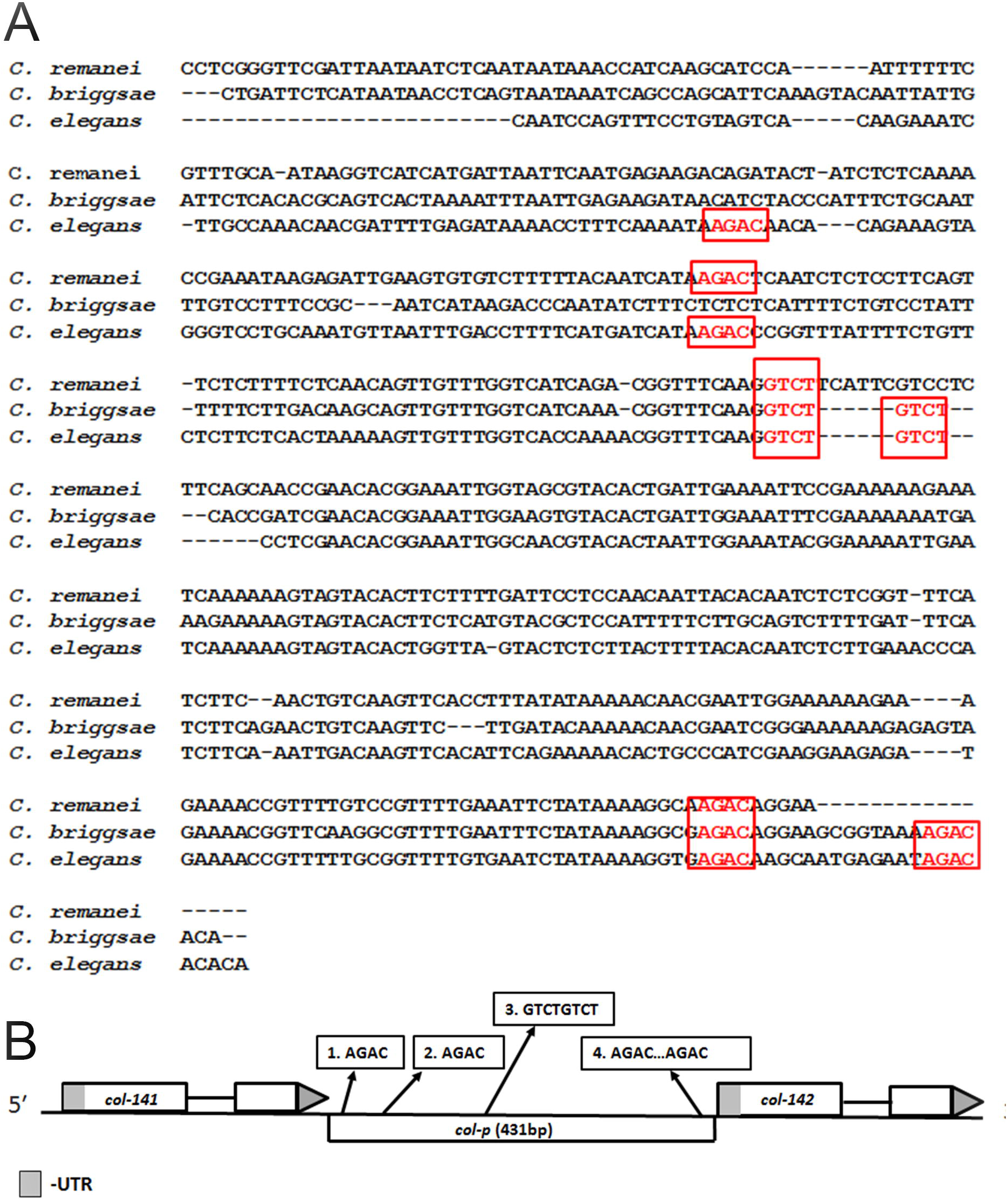
**Multiple conserved SBEs are present in the intergenic region between *col-141* and *col-142*. A.** Five out of the six putative SBE sites are conserved among multiple *Caenorhabditis* species in the intergenic region between *col-141* and *col-142*. Nucleotide alignments were made using ClustalW. **B**. Schematic of the location of SBEs in the intergenic region between *col-141* and *col-142*.

### Cuticle collagen genes may be direct or indirect targets of DBL-1 Smads

In order to elucidate the molecular requirements for Smad recruitment, we used the electrophoretic mobility shift assay (EMSA) to test R-Smad SMA-3 and co-Smad SMA-4 for binding. We generated four probes for EMSA based on sequence alignment and the presence of GTCT sequences (Fig 4C). SMA-4 strongly and specifically bound to probes 3 and 4 (Fig 4A-B). To determine if the conserved SBEs were necessary for SMA-4 binding, we introduced a single base pair substitution in the SBE, changing GTCT to ATCT, which led to loss of specific binding as shown by mutant probes and cold competitor probes. Mutant probes with ATCT are unable to compete with wild type probes for SMA-4 binding (Fig 4B). Our EMSA results indicate that SMA-4 can bind at least two specific sites in *col-p*. Additional binding sites may also be present, because we did not test the entire intergenic region. In contrast to *col-141* and *col-142*, SMA-3 ChIP-seq peaks were not found in the genomic regions surrounding and *col-41*.

**Fig 4.**
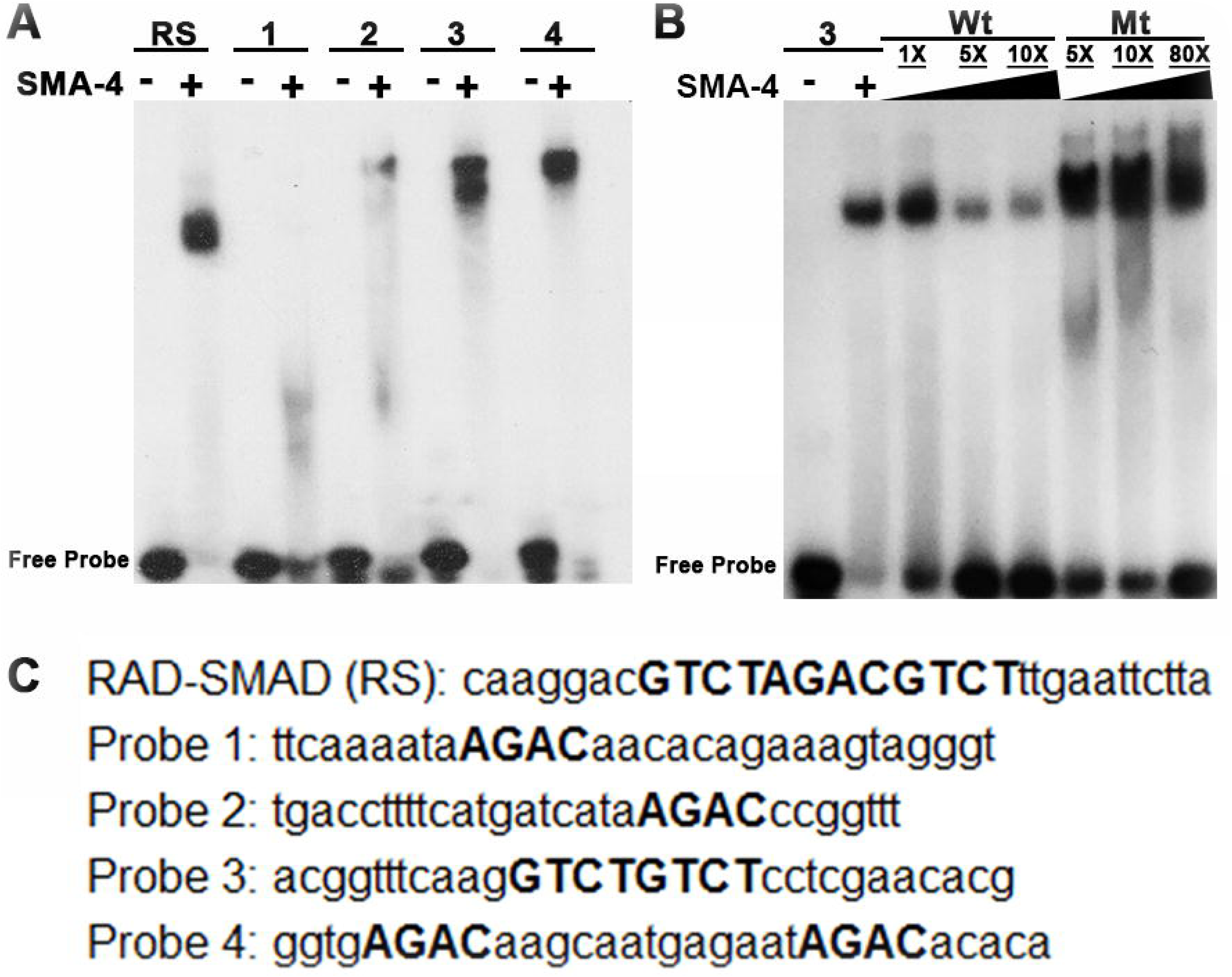
**Smads bind directly to SBEs in *col-p*. A**. SMA-4 strongly binds to probe 3 and probe 4 containing SBEs. **B**. SMA-4 binds specifically to SBE (2xGTCT) in Probe 3. **C**. Probes from *col-p* region used to test binding of Smad MH1 DNA binding domains. SBEs are capitalized in red.

### Identification of genomic sequences required for *col-141* and *col-142* expression

Having identified Smad binding sites in *col-p*, we further aimed to determine whether these sites were relevant for gene expression *in vivo*. Hence, we created a construct with *col-p* driving expression of 2x nuclear localized (NLS) mCherry and obtained multiple lines via microinjection. mCherry expression was observed in the hypodermis and in the vulva at the adult stage (Fig 5A-C), confirming the ability of *col-p* to drive gene expression. Expression was specific to the adult stage only, in alignment with Jackson et al. [25] which showed peak *col-141* and *col-142* expression at the adult stage. To test if the DBL-1 pathway would regulate mCherry expression, we further crossed these lines into *sma-3* mutant background. We observed little to no expression of mCherry expression at all stages (Fig 5D-F). We quantitated fluorescence intensity and found that it was significantly reduced (Fig 5G). This result did not align with our microarray and qRT-PCR results, which showed increased rather than decreased expression of *col-141* and *col-142* in DBL-1 pathway mutants. However, those assays were conducted at the L2 stage, at which time fluorescence of our reporter is undetectable. As described below, qRT-PCR experiments in adult animals are consistent with our reporter results.

**Fig 5.**
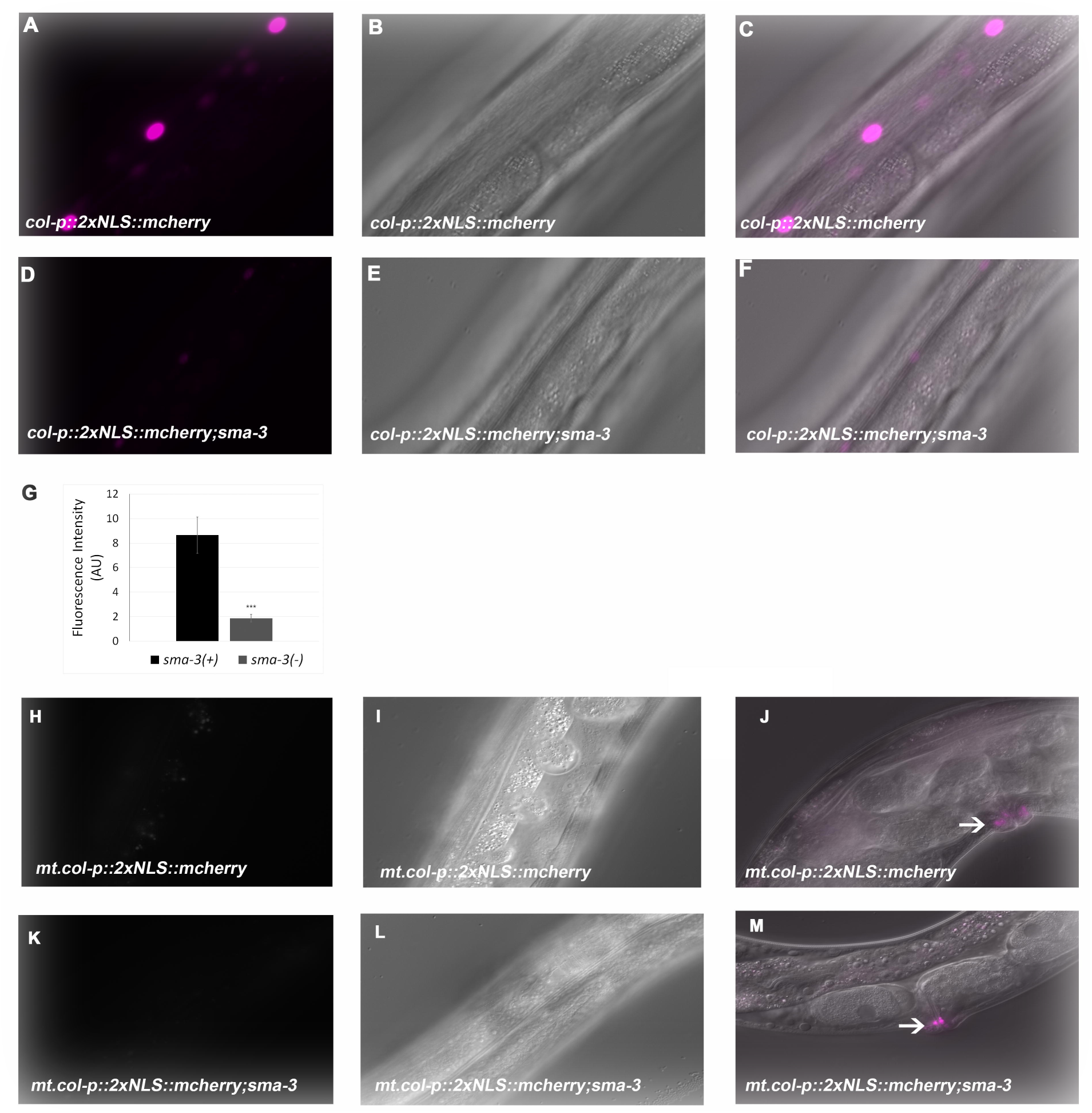
**Identification of genomic sequences required for *col-141* expression. A-C**. *col-p∷2xNLS∷mCherry* leads to adult stage specific expression in the hypodermis nuclei. **D-F**. mCherry expression is severely depleted in *sma-3* mutant background. **G**. Quantification of mCherry expression in wild type and *sma-3* mutant background. Error bars show standard error. * P < 0.05, ** P < 0.005, *** P < 0.0005 **H-J**. Mutating SBE 3 from GTCT to ATCA, SBE 4 from AGAC to TGAT leads to loss of mCherry expression. **K-M**. Crossing of *mt.col-p∷2xNLS∷mCherry* into *sma-3* mutant background does not change the ablated mCherry expression. **A,D,H,K** fluorescent images; **B,E,I,L** Nomarski images; **C,F** merged images; **J,M** merged images showing expression in the vulval region.

To test if SBEs present in *col-p* are relevant for mCherry expression, we made transgenic animals with mutated SBEs (*mt.col-p)* driving 2xNLS∷mCherry. We noted complete loss of mCherry expression except in the vulva (Fig 5J), indicating Smad binding at these SBEs is required for gene expression in the hypodermis (Fig 5H-I). In order to test if mCherry expression was the same in the presence and absence of *sma-3,* we crossed these worms into *sma-3* mutant background (Fig 5K-M). Again, we observed a complete absence of fluorescence.

To verify the results from our extrachromosomal reporter, we wanted to study the expression of *col-141* in its endogenous genomic environment. Utilizing CRISPR-Cas9, we knocked in 2xNLS∷GFP to replace *col-141* making a transcriptional reporter and a loss of function simultaneously (Fig 6A). We characterized the reporter and observed that GFP expression is only visible at the adult stage in the hypodermis alone (Fig 6B-D), with little to no expression observed in larval stages, consistent with the result from the extrachromosomal reporter. To confirm whether DBL-1 signaling transcriptionally regulated GFP expression, we crossed the *col-141* reporter into a *sma-3* mutant background (Fig 6E-G). We observed a loss of GFP expression, replicating the results of the extrachromosomal *col-p∷2xNLSmCherry* transgenic animals (Fig 5A,D).

**Fig 6.**
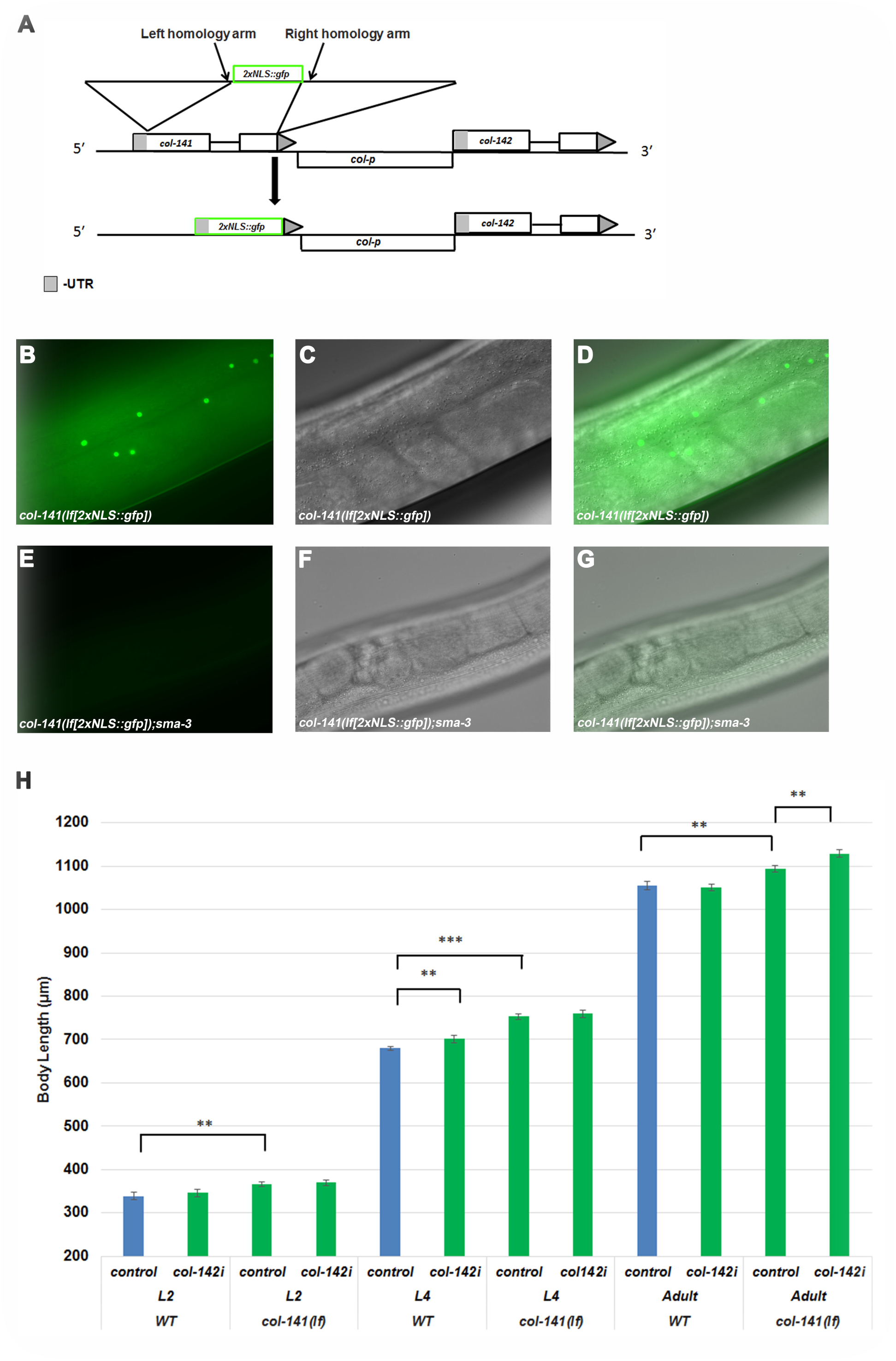
**An endogenous CRISPR-generated reporter verifies requirements for *col-141* expression. A.** Utilizing CRISPR-Cas9 genome editing to knock out and replace *col-141* with 2xNLS∷gfp, creating a loss of function mutant and transcriptional reporter simultaneously. **B-D**. GFP expression in *col-141(lf)* mutant. **E-G**. Loss of GFP expression in *sma-3* mutant background. **B,E** fluorescent images; **C,F** Nomarski images; **D,G** merged images. H. Body size phenotypes of *col-141(lf), col-142(RNAi)* and doubles. Error bars show standard error. * P < 0.05, ** P < 0.005, *** P < 0.0005.

This genome-edited animal also allowed us to determine the body size phenotype of *col-141(lf)* and compare it to *col-141(*RNAi). We measured body length of *col-141(lf)* at L2, L4, and adult stages and found that it is longer than controls at all three stages (Fig 5H, S2), similar to the RNAi results (Fig 1A) except that body length differences persisted into adulthood in the mutants. Since *col-141* shares sequence homology with *col-142*, we wondered if there could be partial functional redundancy between the two genes. We therefore performed RNAi targeting *col-142* on the *col-141(lf)* mutant and observed a further increase in body size in adults compared to *col-141(lf)* alone, and compared to wild type (Fig 5H).

### DBL-1 signaling is required for stage-specific regulation of cuticle collagen genes

We sought to resolve our discrepant results on the direction of regulation of *col-141* and *col-142* by the DBL-1 pathway. Initial qRT-PCR performed at the L2 stage indicated an increase in *col-141* and *col-142* expression in DBL-1 pathway mutants (Table 1). Meanwhile, the *col-p∷2xNLSmCherry* and *col-141(lf[2xNLS::gfp])* strains showed a reduction of expression at the adult stage. In order to confirm that loss of fluorescence expression in *col-141(lf[2xNLS::gfp])* and *col-p::2xNLSmCherry* replicated *col-141* and *col-142* expression in *sma-3* mutant background, we performed qRT-PCR on *sma-3* mutant adults. We observed a significant decrease in *col-141* and *col-142* levels in *sma-3* mutant adults (Table 1), corroborating the observations from our transcriptional reporters. Therefore, the direction of action of the DBL-1 pathway on *col-141* and *col-142* expression is stage-specific. We extended these observations by determining the expression levels of *col-41* and *rol-6* in *sma-3* mutant adults. In these experiments, we observed an increase in *col-41* and *rol-6* levels (Table 1), again reversing the direct of regulation in comparison with the L2 stage. These results suggest that DBL-1 signaling is required for the appropriate temporal stage-specific expression of cuticle collagen genes.

## Discussion

Body size regulation has been investigated in multiple organisms, including humans [33, 34] mice, dogs, and Drosophila [35–38]. In these organisms, signaling pathways have been identified that contribute to body size regulation; however, the effector genes for body size regulation through those pathways are poorly characterized. We have capitalized on the genetic tractability of *C. elegans* to study how the conserved DBL-1 BMP-related signaling pathway regulates body size. In our work, we elucidated the transcriptional network initiated by DBL-1 to explain how body size is modulated by DBL-1 activity. We have shown that cuticle collagen genes are transcriptional targets of the DBL-1 pathway via microarray analysis and qRT-PCR. Moreover, we showed that DBL-1 is required for stage-specific expression of cuticle collagen genes. Through RNAi, mutant, and overexpression studies we discovered a positive regulator (*col-41*), a dose-sensitive regulator (*rol-6*) and negative regulators (*col-141, col-142*) of body size. We showed association of Smads in the intergenic region between *col-141* and *col-142* via ChIP-seq and using EMSA we showed *in vitro* binding of SMA-4 MH1 domain to conserved SBE sites in this region. Additionally, we show lack of binding by Smads to the previously defined *col-41* promoter region. Hence, cuticle collagen genes are directly and indirectly regulated via the DBL-1 pathway. Based on presented evidence, we conclude that cuticle collagen genes are major effectors of body size through the DBL-1 pathway. We speculate that the specific collagen isoforms deposited in the cuticle contribute to the characteristics of that cuticle, due to differences in how they crosslink with other components. COL-41, as a positive regulator of growth, may contribute to a more elastic, expandable cuticle. COL-141 and COL-142, as negative regulators of growth, may contribute to a more rigid, less easily expanded cuticle.

Since cuticle collagen genes are encoded by such a large gene family in *C. elegans,* it is reasonable to ask why so many genes are needed. One possible explanation is that the large number of genes allows rapid synthesis of the cuticle during the discrete cuticle synthesis periods of the worm’s lifecycle. Our work supports the notion that there is also a degree of functional specificity. Consistent with this hypothesis, more than 60% of cuticle collagen genes have been described as having stage-specific expression patterns [25]. In particular, *col-41* and *rol-6* have peaks of expression in the L2 stage, while *col-141* and *col-142* have highest expression in adults. In addition to our own work, two published microarray analyses identified cuticle collagen genes as transcriptional targets of the DBL-1 pathway [23, 28, 39]. Furthermore, specific subsets of cuticle collagen genes have been described as transcriptional targets of Wnt and insulin signaling pathways in *C. elegans* [25, 40]. Both DBL-1 and Wnt pathway mutants have been shown to have cuticle defects [25, 41], demonstrating that there are functional consequences to the alterations in collagen gene expression. Furthermore, Ewald et al. [40] demonstrated that specific cuticle collagens (which include *col-141*) contribute to the regulation of longevity by the insulin signaling pathway.

In humans, collagens constitute ~30% of the protein mass and are the most abundant protein. Collagens function in a diverse range of processes such as mechanical properties of tissue, ligands for receptors, cell growth, differentiation, cell migration, bone formation, skeletogenesis, and the integrity and function of the epidermis [42–44]. The expression levels of collagens must be tightly controlled during wound healing, and inappropriate expression of collagens can lead to fibrosis [45, 46]. Mutations in collagens have been recognized in rare heritable diseases such as osteogenesis imperfecta, several subtypes of Ehlers-Danlos syndrome, various chondrodysplasias, X-linked forms of Alpoty syndrome, Ullrich muscular dystrophy, corneal endothelial dystrophy, and Knobloch syndrome [47]. Therefore, collagens and their regulation have significant implications for human health. Our work further adds to the functional importance of collagens in the context of growth regulation.

In addition to highlighting the functional significance of collagen genes, our work sheds light on the mechanisms of action of the DBL-1 Smads. Three Smads act in this pathway: the R-Smads SMA-2 and SMA-3 and the co-Smad SMA-4. All three Smads are necessary for pathway function, and they presumably form a heterotrimeric complex. We performed genome-wide ChIP-seq analysis on GFP∷SMA-3 and identified GTCT, the canonical SBE, as enriched at sites of SMA-3 occupancy. In Drosophila, Mad, the Dpp R-Smad, binds GC-rich Smad binding sites termed GC-SBE (GGAGCC) [48], while Medea, the Dpp co-Smad, binds canonical SBEs (GTCT). In our analyses of targets of the DBL-1 pathway, we observed Smad binding to the intergenic region between *col-141* and *col-142*. We therefore tested for direct binding of isolated MH1 domains of Smads SMA-3 and SMA-4 to SBEs in this region. Consistent with findings in Drosophila, the MH1 domain of co-Smad SMA-4, but not that of R-Smads SMA-3, was able to bind the SBE’s directly *in vitro.* Similarly, the co-Smad of the *C. elegans* DAF-7/TGF-β pathway, DAF-3, has previously been shown to bind canonical GTCT sequences [49]. Since there is no evidence for direct binding of R-Smad SMA-3 to GTCT sequences, the enrichment of this motif at SMA-3 genomic binding sites may be due to SMA-3 being recruited as part of the Smad complex, with co-Smad SMA-4 making the direct contact with this site.

In conclusion, our work elucidates the transcriptional network for body size regulation through the DBL-1/BMP pathway in *C. elegans.* We propose a model (Fig 7) in which DBL-1 signaling leads to Smad activation causing direct and indirect regulation of specific cuticle collagen genes. This regulation occurs in a stage-specific manner to initiate the correct temporal program of collagen gene expression. Inappropriate expression of cuticle collagen genes leads to aberrant body size. This work thus provides the first mechanistic link between BMP signaling and effectors of body size regulation.

**Fig 7.**
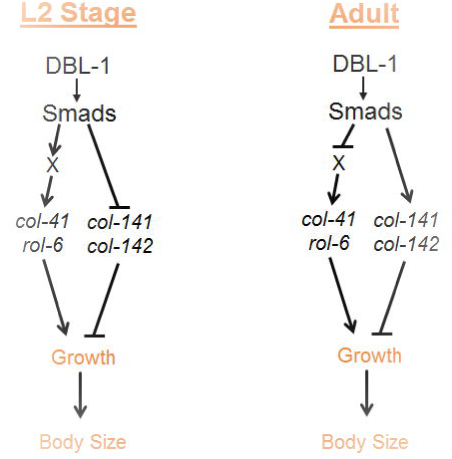
**DBL-1 regulation of growth.** Our model proposes stage-specific regulation of cuticle collagen genes to contribute to growth and body size of *C. elegans*.

## Materials and Methods

### RNA interference (RNAi)

RNAi by feeding was performed as described [50, 51]. RNAi clones that target the unique (non-Gly-X-Y repeat) regions of collagen genes were generated by cloning desired PCR fragments into L4440.

### Electrophoretic Mobility Shift Assay (EMSA)

*Protein purification:* GST tagged versions of the MH1 domains of SMA-2, SMA-3, and SMA-4 were cloned in the pGEX-4T2 vector. All three proteins were isolated using BL21 competent cells, grown in a starter culture O/N at 37°*C*. From the starter culture a fresh culture of 100 ml was started, grown at 37°*C* until O.D. of 500 was reached. Expression was induced using IPTG at final concentration of 1 mM in 100 ml cultures and grown at 28°C for 6-8 hours. Standard batch purification was performed using GST beads and eluted in 30 mM reduced glutathione and 50 mM Tris buffer, pH 8.0. Three elutions were taken, each of the primary elutions were run on 4-12% Bis-Tris gels (Life Technology) for size separation under denaturing conditions. EMSA *Gel Shifts:* Double stranded probes were labeled with ^32^P and incubated with protein in binding buffer at RT for 20 minutes. Binding Buffer was composed of 50% glycerol, 100mM HEPES, 15mM DTT, 0.5mg/ml BSA, 0.5M KCl, 50mM ZnSO_4,_ 50 μg/ml dIdC, samples were run on 5% native acrylamide gel for 1-1.5 hours, dried for one hour at 65°C and developed for 1 hour at −80°C. Probe 1: ttcaaaataagacaacacagaaagtagggt. Probe 2: tgaccttttcatgatcataagacccggttt. Probe 3: acggtttcaaggtctgtctcctcgaacacg. Probe 4: ggtgagacaagcaatgagaatagacacaca.

### qRT-PCR

Worms were grown at 20°C until a large amount of eggs were observed on plates. Worms were then washed off using M9 buffer and the remaining eggs were allowed to hatch for 4 hours. Worms were then collected and placed on new plates and grown at 20°C until late L2 or adult stage, collected and mRNA was extracted using the RNeasy mini kit (Qiagen). Reverse transcriptase PCR and quantitative real time PCR were performed as previously described [52].

### ChIP-Seq

ChIP-seq was performed by Michelle Kudron (Valerie Reinke modENCODE/modERN group) on *sma-3(wk30); Is*[GFP∷SMA-3] strain at late L2 stage [53].

### Genome editing and transgenic strains

To obtain a *col-141* knock-in line via CRISPR-Cas9 we followed the Dickinson [54] protocol and used the selection excision cassette pDD282 vector. All constructs were assembled using NEB Hi-fi DNA Assembly kit. All other strains were constructed by microinjections, *myo-2∷gfp* (20 ng/μl), transgene of interest (100 ng/μl), carrier DNA (80 ng/μl). *col-141* and *col-142* overexpression lines were created via microinjection (50 ng/μl each, *myo-2: gfp* at 12 ng/μl). *rol-6* overexpression lines were made via injection of *rol-6* (50 ng/μl) and *myo-2∷gfp* (10 ng/μl).

## Acknowledgements

We thank Michelle Kudron with the modENCODE and modERN projects for performing ChIP-seq; Randy Hui for technical assistance with EMSA; Amy Park for RNAi experiments; and Emma Ciccarelli for constructing the *col-142* RNAi clone. We are grateful to Jun (Kelly) Liu and Geraldine Seydoux for sharing protocols and Hannes Buelow for providing clones. Some strains were provided by the CGC, which is funded by NIH Office of Research Infrastructure Programs (P40OD010440).

---

**Fig S1. Body size following RNAi of DBL-1 transcriptional targets that encode potential endoreduplication regulators.** Body length was measured at 96 hours after embryo collection. Error bars show standard error.

**Fig S2. Body size of *col-141(lf)* CRISPR strain.** Body size was measured at L2, L4 and adult stages. Error bars show standard error. *** P < 0.0005.

